# The immune checkpoint kick start: Optimization of neoadjuvant combination therapy using game theory

**DOI:** 10.1101/349142

**Authors:** Jeffrey West, Mark Robertson-Tessi, Kimberly Luddy, Derek S. Park, Drew F.K. Williamson, Cathal Harmon, Hung T. Khong, Joel Brown, Alexander R.A. Anderson

## Abstract

An upcoming clinical trial at the Moffitt Cancer Center for women with stage 2/3 ER+breast cancer combines an aromatase inhibitor and a PD-L1 checkpoint inhibitor, and aims to lower a preoperative endocrine prognostic index (PEPI) that correlates with relapse-free survival. PEPI is fundamentally a static index, measured at the end of neoadjuvant therapy before surgery. We develop a mathematical model of the essential components of the PEPI score in order to identify successful combination therapy regimens that minimize both tumor burden and metastatic potential, based on time-dependent trade-offs in the system. We consider two molecular traits, CCR7 and PD-L1 which correlate with treatment response and increased metastatic risk. We use a matrix game model with the four phenotypic strategies to examine the frequency-dependent interactions of cancer cells. This game was embedded into an ecological model of tumor population growth dynamics. The resulting model predicts both evolutionary and ecological dynamics that track with changes in the PEPI score. We consider various treatment regimens based on combinations of the two therapies with drug holidays. By considering the trade off between tumor burden and metastatic potential, the optimal therapy plan was found to be a 1 month kick start of the immune checkpoint inhibitor followed by five months of continuous combination therapy. Relative to a protocol with both therapeutics given together from the start, this delayed regimen results in transient sub-optimal tumor regression while maintaining a phenotypic constitution that is more amenable to fast tumor regression for the final five months of therapy. The mathematical model provides a useful abstraction of clinical intuition, enabling hypothesis generation and testing of clinical assumptions.

## 1. Introduction

Evolving alongside vacillating populations of pathogens, the components of human immune system mirror the changing and hostile environments of our past. While capable of defending against pathogens and dysfunctional cells, selection events result in trade-offs between protective responses, autoimmunity, and host fecundity, shaping the diverse cellular and molecular composition of the immune system. Immune regulation serves to protect healthy tissue during a powerful immune response. In tumors, cancer cells often evolve strategies to dysregulate, coopt or suppress the immune system. Recently, therapies that block such immune evasion mechanisms, using antibodies that target immune checkpoints (e.g. cytotoxic T lymphocyte antigen-4 (CTLA-4)and programmed death-1 (PD-1)), have been introduced to enhance anti-tumor cytotoxic T cell responses by inhibiting immune regulatory functions [1]. The efficacy of immune checkpoint blockade (ICB) therapy was first confirmed in overall survival of advanced melanoma patients treated with the anti-CTLA-4 monoclonal antibody ipilimumab [2, 3]. Subsequently anti-PD-L1 therapy was included as a broadly applicable tool for the treatment of cancer [4, 5, 6]. PD-1 is a checkpoint protein expressed by cytotoxic immune cells (T cells and NK cells) and its ligand PD-L1 is often expressed on cancer cells, leading to immune evasion. Using ICB to block either CTLA-4 or PD-L1 has led to durable responses in some patients, albeit with only a fraction of patients responding, possibly due to other immune evasion mechanisms [6, 7]. Therefore, therapeutic approaches using combinations of ICB and targeted therapies or radiotherapies, that stimulate various steps of the cancer-immunity cycle, need to be explored.

### 1.1. Combination therapy with immune checkpoint inhibitors

Chemotherapy and radiation are widely used and readily combined with ICB therapies, reviewed in detail in Patel & Minn [7]. Radiation improves responses to ICB in mice given anti-CTLA-4 [8, 9]. There also seems to be an additional abscopal effect (tumor shrinkage in unirradiated areas) [10]. Mathematical models illustrate the potential synergies with immune checkpoint inhibitors [11, 12]. Similarly, genotoxic chemotherapies combine favorably with ICB therapies by improving the discriminatory function of the immune system to evoke immunogenic pattern recognition receptor (PRR) signalling [13, 14]. Previously, Radfar et al. conducted a study of an approach termed “chemocentric chemoimmunotherapy with potential application in the treatment of all cancer types [15]. The technique utilizes activated CD4^+^ T cells to chemosensitize the tumor before chemotherapy administration. These results support other recent studies reporting improved response rates and survival with salvage chemotherapy in patients who previously received cancer vaccination aimed at developing an immune response [16, 17, 18, 19, 20].

Further, multiple classes of targeted therapies in combination with ICB show promising results, for example CDK4/6 inhibitors used in hormone receptor-positive breast cancer [21, 22, 23]. Pre-clinical experimental data shows synergism between targeted therapy (a BRAF inhibitor) and anti-PD-L1 therapies that counteract subsequent immune escape via expression of PD-L1 that would typically occur in tumors of patients with metastatic melanoma [24, 25].

There are currently over 400 ongoing trials of ICB in combination with other modalities. While combination therapies have tremendous clinical potential, designing combination trials requires a deep understanding of the underlying biological mechanisms at play in two complex and interacting systems: the tumor and the immune system. Challenges magnify when confronting the myriad of possible dose levels and timing schedules. The clinical options become legion, even though they all have the same goal of eradicating the tumor while preventing the emergence of resistance. Mathematical modelling permits exploration of the myriad of options as well as integrating knowledge and assumptions from the clinic and research bench. The goal of our modelling here is to generate testable hypotheses [26]. Previously, mathematical models have been used to study tumor-immune dynamics and system stability in a variety of cancer types [27, 28, 29, 30, 31].

### 1.2. Neoadjuvant therapy in ER^+^ breast cancer

A recently approved clinical trial, at Moffitt, will treat women with stage 2 or 3 breast cancer. All patients will have estrogen-receptor positive (ER^+^) tumors and receive neoadjuvant combination therapy: an aromatase inhibitor (primarily anastrozole) and checkpoint inhibitor against PD-L1 (durvalumab). Aromatase inhibitors (AI) are an orally available class of drugs used in post-menopausal ER^+^ breast cancer patients. AI therapy suppresses estrogen production by blocking the aromatase enzyme, a key step in estrogen production. ER^+^ breast cancer cells depend on estrogen for proliferation. By reducing the amount of hormone available to tumor cells, AI therapy blocks the growth of the tumor resulting in tumor cell death. Response rates to AI therapy are relatively high (50% to 70%) in the neoadjuvant setting but there still remains a large population of patients, especially with advanced disease, that do not respond. There is a need for alternative therapies, as well as biomarkers to predict response [32]. The addition of checkpoint inhibitors to AI drug regimens appears promising [33, 34, 35]. Therefore a key goal of this paper is to provide insights into how best to deliver this combination therapy.

### 1.3. Mathematical modeling of immune checkpoint inhibitors

Here we develop a mathematical framework to better understand the effect of combining AI therapy with the checkpoint inhibitor (CI) anti-PD-L1 in post-menopausal ER^+^ breast cancer. Specifically, we base our model on the clinical trial discussed above, which begins accruing patients in mid-2018. The trial has a short-term outcome metric, the preoperative endocrine prognostic index (PEPI) measured at the end of six months of neoadjuvant therapy [36]. The PEPI score accounts for tumor size, presence of cancer cells in the lymph nodes, Ki-67 levels (a marker for proliferation), and hormone receptor status. The PEPI score provides a prognostic indicator for relapse-free survival (RFS). The PEPI score is measured at a single time point, namely at the time of surgery, following neoadjuvant therapy. The mathematical model developed here mimics two essential indicators of the PEPI score, namely lymphatic metastatic potential, and tumor size. We use an evolutionary game theory (EGT) approach in order to 1) evaluate the success of diverse combination therapies of AI and CI that aim to minimize both tumor burden and metastatic potential; and 2) identify the *time-dependent* trade-offs between tumor burden and metastatic potential within the PEPI score.

We consider CCR7 and PD-L1 as two relevant and measurable molecular traits of the cancer cell that play a role in therapy resistance. Furthermore, both PD-L1 and CCR7 are positively correlated with metastatic risk, especially to the lymphatics [37, 38, 39]. Similar to the PD-1/PD-L1 axis, CC-chemokine receptor 7 (CCR7) and its ligands (CCL19/21) help to govern the balance between immune activation and immune regulation. Lymphocyte trafficking is a highly regulated process guided by chemokine gradients and integrins. CCR7 is a chemokine receptor traditionally expressed by immune cells and is required for leukocyte homing to lymph-nodes. Along with other chemokine receptors, CCR7 expression in cancer cells is highly correlated with lymph node involvement and metastasis in breast cancer [37, 38, 39]. Additionally, CCR7 activation induces proliferation and inhibits apoptosis in both immune cells and cancer cells [40, 41, 42]. In a small study examining gene expression in 89 patients receiving letrozol, CCR7 expression was elevated in nonresponders compared to responders. As seen in Figure 1, CCR7 expression increases in all patients (light grey) and to a greater degree in nonresponders (dark grey) [32]. Here, we employ CCR7 expression as a marker of metastatic potential with increased risk of lymph-node invasion. Furthermore, we see positive CCR7 expression as conferring resistance to the aromatase inhibitor.

**Figure 1:**
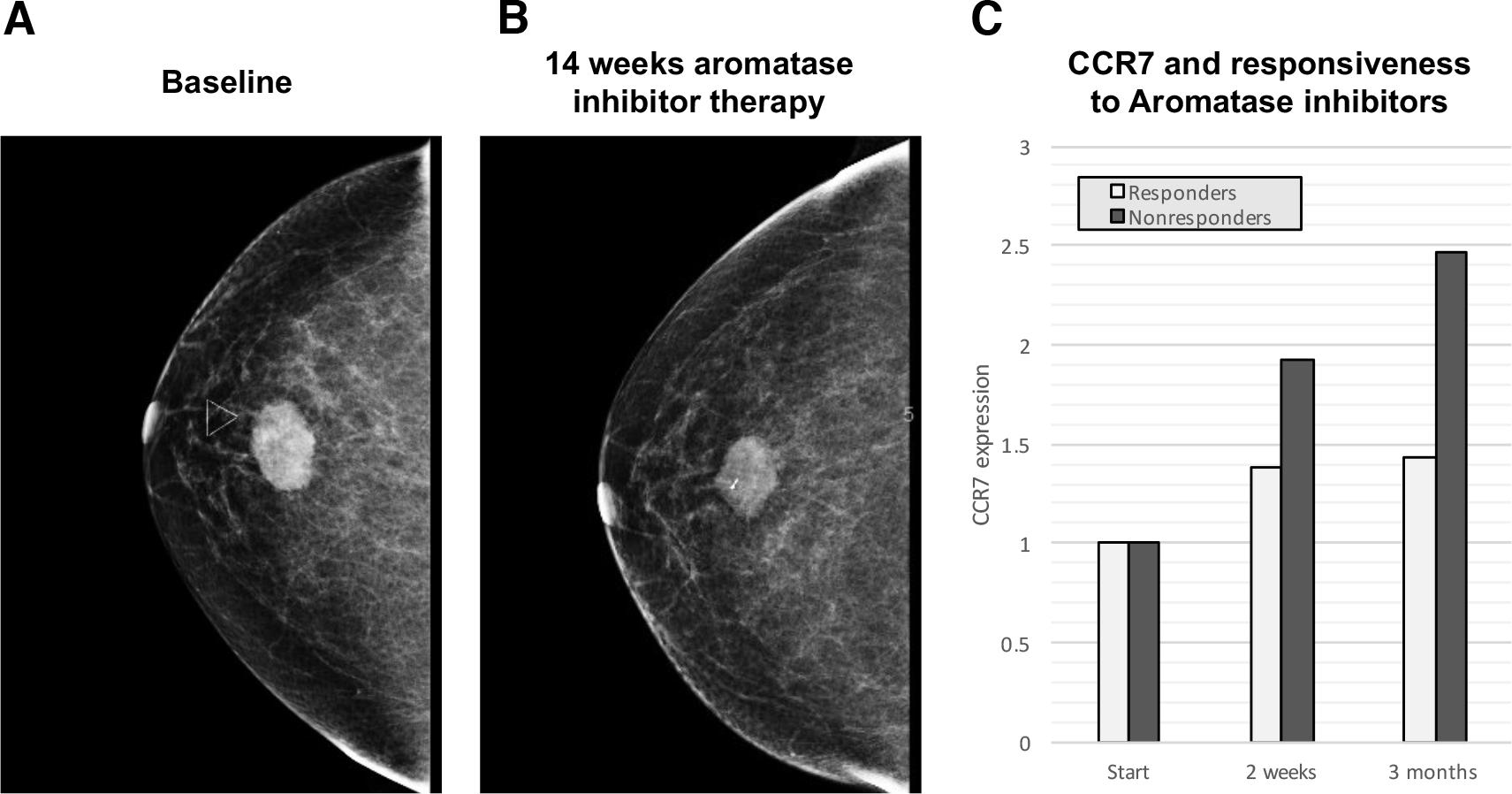
Expression changes during continuous administration of aromatase inhibitor therapies. — (a,b) A representative image before (a) and after (b) neoadjuvant aromatase inhibitor therapy shows a non-responder (30% to 50% of patients) to the current standard of care. (c) Expression data from biopsies obtained from 89 postmenopausal women with ER^+^ breast cancer receiving neoadjuvant letrozole [32] were analyzed. CCR7 was identified in a group of genes which showed the greatest expression pattern changes during treatment with AI. Gene ontology analysis via AmiGO (amigo.geneontology.org) revealed that this subset of differentially expressed genes was enriched for immune function, including CCR7 as a marker of potential metastatic escape. Relative changes of CCR7 expression increased under AI therapy in all patients, but to a greater degree for nonresponders.

By using CCR7 and PD-L1 tumor phenotypes in the mathematical model, we incorporate key elements of therapy resistance strategies that go on to influence the PEPI score (tumor size and risk of lymph metastasis). The model provides the evolutionary trajectories of these cancer cell phenotypes and how they impact the total cancer cell population over time. In order to combine the evolutionary dynamics with the population dynamics we embed an evolutionary game theory model of cell-type specific interactions into a model of population dynamics. Thus, we integrate the replicator dynamics (changes in strategy frequencies) with the population dynamics. The model serves as a platform to aid clinical intuition in designing combinations of hormone therapy and immunotherapy.

## 2. Methods

Evolutionary game theory provides a framework for modelling frequency dependent selection where the fitness value of a trait to an individual depends on the trait values of others. A payoff matrix defines the fitness returns to an individual (row strategy) from interacting with another individual or the population at large (column strategy) [43, 44, 45]. It is now well established that cancer progression is an evolutionary and ecological process [46, 47] where evolutionary forces (such as genetic drift with heritable mutations and natural selection) drive changes in the cancer cells’ heritable phenotypes along a fitness landscape [48, 49]. Evolutionary game theory has previously been used for modelling cancer treatment, including models of prostate cancer tumor-stroma interactions [50, 51], adaptive therapy [52], metronomic chemotherapy [53], competitive release [54] and the evolutionary double-bind [55].

Our EGT model sees tumor cells as “players” with two independent phenotypic axes: CCR7 and PD-L1 expression. Each cell of type *i* (*i* = 1, 2, 3, 4) competes according to equation (1), where 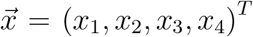 is the vector of the corresponding frequency of the 4 phenotypes: *x*_1_ ≡ CCR7^−^ / PD-L1^−^; *x*_2_ ≡ CCR7^−^ / PD-L1^+^; *x*_3_ ≡ CCR7^+^ / PD-L1^−^; and *x*_4_ ≡ CCR7^+^ / PD-L1^+^. The fraction of cells expressing CCR7 and PD-L1 can be found by *x*_3_ + *x*_4_ and *x*_2_ + *x*_4_, respectively. The evolution of average trait values within the tumor population can be tracked on a two-dimensional plane (Figure 2A).

**Figure 2:**
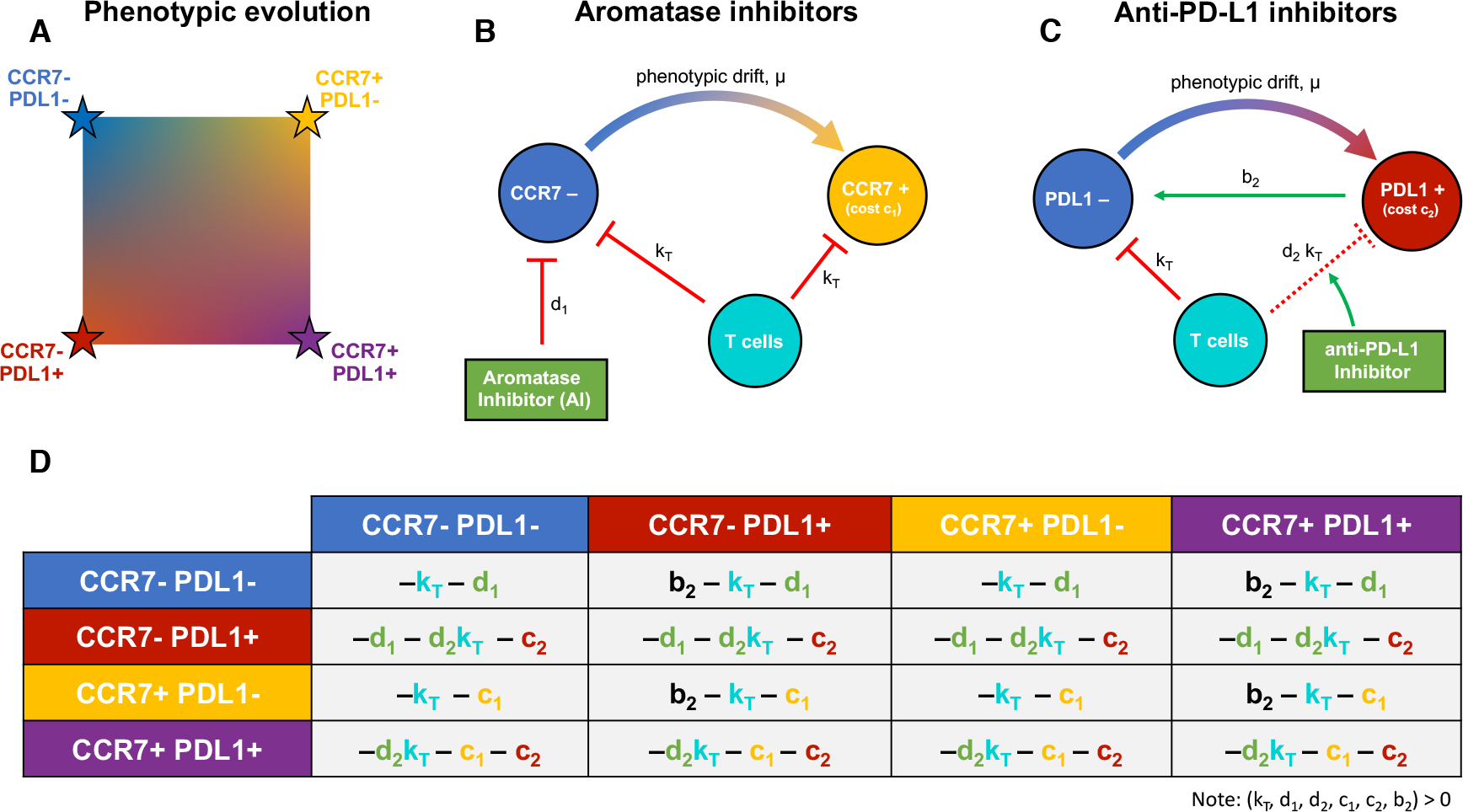
Schematic of the Mathematical Model. — (a) The tumor population consists of phenotypes varying on two axes: CCR7 (increasing from left to right) and PD-L1 (increasing from top to bottom). (b) AI therapy targets CCR7^−^ phenotypes with kill rate of *d*_1_; these may evolve CCR7 expression due to selection. T cells kill both types at rate *k*_*T*_. (c) CI therapy boosts the immune response against tumor cells expressing PD-L1. Additionally, PD-L1^−^ cells benefit from being near PD-L1^+^ cells, known as a “cheater” population. (d) The two-dimensional phenotypic space is modeled using a four player game with subscript 1 indicating the CCR7 axis and subscript 2 indicating the PD-L1 axis. Killing terms (see red lines in b,c) for *k*_*T*_, *d*_1_, *d*_2_ are added to PD-L1^−^, CCR7^−^, and PD-L1^+^ rows respectively. Cost terms (*c*_1_, *c*_2_) are added to CCR7^+^ and PD-L1^+^ rows respectively. The “cheater” benefit *b*_2_ is added to PD-L1^−^ interactions (row) with PD-L1^+^ cells (column). Note: all costs (*c*_1_, *c*_2_, *k*_*T*_, *d*_1_, *d*_2_) and benefits (*b*_2_) in payoff matrix *A* are constrained to non-negative values.

The phenotypic interactions for each therapy are shown in Figure 2B and 2C. T cells kill tumor cells at rate *k*_*T*_, but PD-L1 expression removes this term and provides a benefit *b*_2_ to neighboring PD-L1^−^ cells. AI targets CCR7^−^ cells (*d*_1_) and CI promotes T-cell killing of PD-L1^+^ cells (*d*_2_*k*_*T*_).

The costs and benefits attributed to each phenotype are mathematically represented in a “payoff matrix” (Figure 2D). The fitness of each phenotype subpopulation is given by equation 3: a function of phenotype prevalence weighted by the payoff matrix. The subscript 1 is associated with AI therapy while subscript 2 is associated with CI therapy, and the parameters are summarized as follows:

- *k*_*T*_: immune kill – the rate at which T cells kill cancer cells; *k*_*T*_ ≥ 0
- *d*_1_: AI drug kill rate; *d*_1_ ≥ 0
- *d*_2_: CI binding blocking rate; 1 ≥ *d*_2_ ≥ 0
- *c*_1_: inherent cost of developing increased CCR7 expression; *c*_1_ ≥ 0
- *c*_2_: inherent cost of developing PD-L1 expression; *c*_2_ ≥ 0
- *b*_2_: “cheater” benefit that a PD-L1^−^ cell gains by being near a PD-L1^+^ cell; *b*_2_ ≥ 0

The total volume, *υ*_*i*_(*t*) for each *i*th phenotype increases by growth rate, *λ*, offset by the various costs and benefits in the fitness function. In this way, the *population* dynamics (*υ*_*i*_(*t*)) are dependent on *absolute fitness*, while the *evolutionary* dynamics (*x*_*i*_(*t*)) are dependent on *relative fitness*. The total change in volume of the tumor is given by the sum of phenotype volume changes: 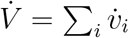.

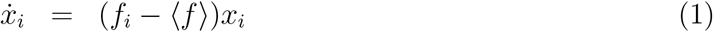

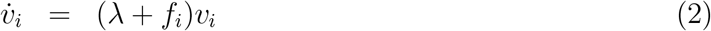

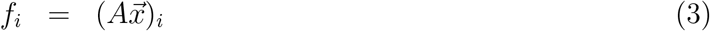

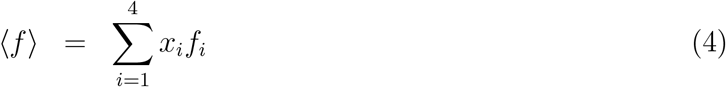

## 3. Results

Figure 3 shows the model predicted treatment responses to: 1) no treatment (black, exponential growth), 2) AI continuous treatment only (blue), 3) CI continuous treatment only (red), 4) both AI and CI continuous treatment (purple). Immune kill is always present (*k*_*T*_ = 0.4) but is not sufficient to cause tumor regression by itself. The costs to develop resistance are small (*c*_1_ = *c*_2_ = 0.1), as is the “cheater” benefit that a PD-L1^−^ cell derives from being near a PD-L1^+^ cell, chosen as *b*_2_ = 0.1. The costs to both CCR7 and PD-L1 expression outweigh the cheating benefit of residing near a PD-L1^+^ cell in the presence of AI drug kill: (−(*c*_1_ + *c*_2_) > *b*_2_ + *d*_2_*k_T_* − *k*_*T*_) − *d*_1_). We assume a low CCR7 expression (*x*_3_ + *x*_4_) and PD-L1 expression (*x*_2_ + *x*_4_) at the start of treatment. We use the simplifying assumption that 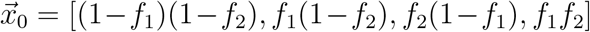 where *f*_1_ the initial fraction of CCR7 expression (*x*_3_ + *x*_4_) and *f*_2_ is initial fraction of PD-L1 expression (*x*_2_ + *x*_4_).

**Figure 3:**
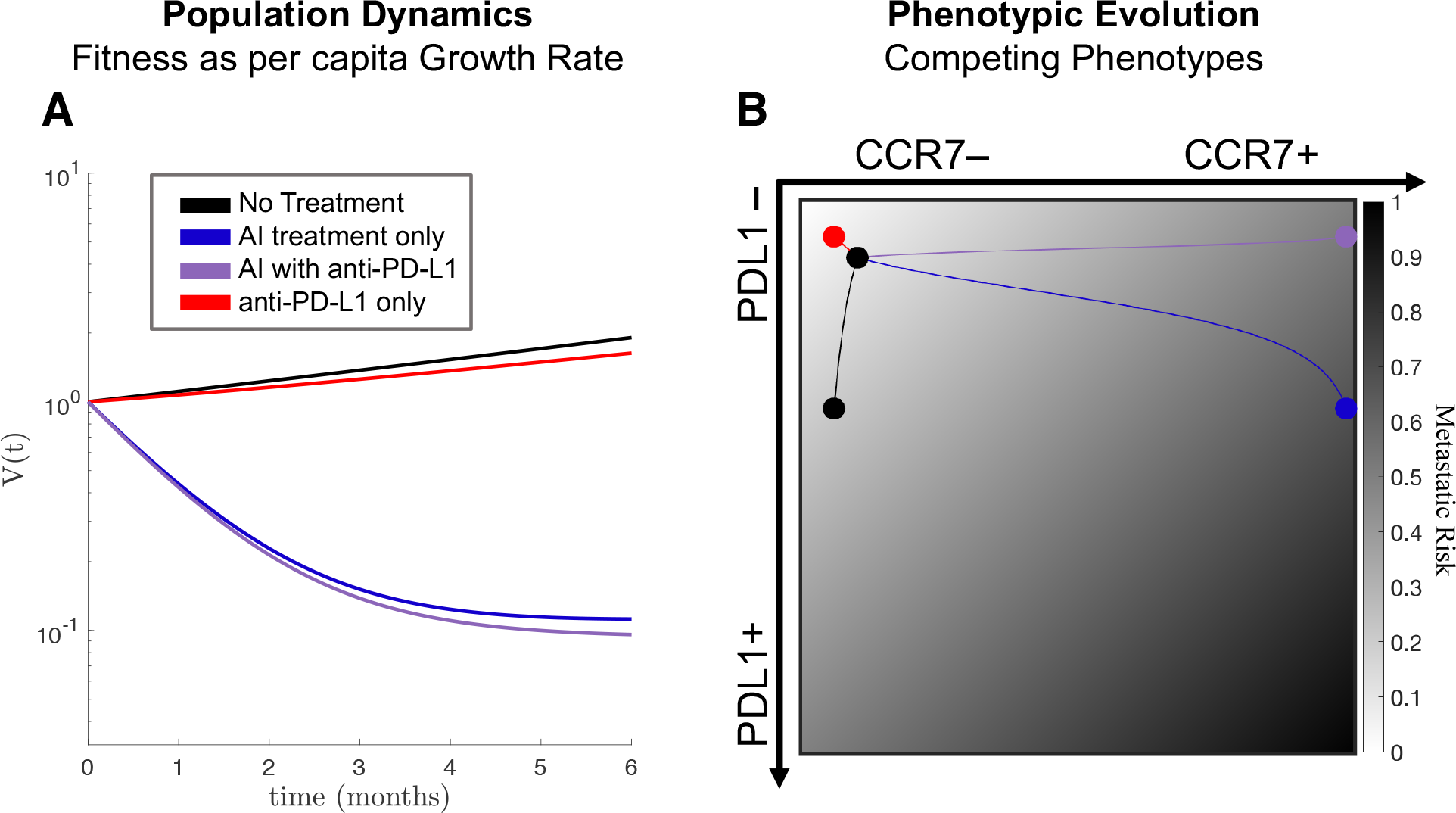
Trade-off between metastatic risk and tumor regression. — (a) Trajectories on the metastatic risk simplex (white to black for increased risk) are shown for no treatment (red), continuous AI (blue), continuous CI (yellow) and combined therapy (green). (b) Tumor regression is shown for the four scenarios. Treatments may be associated with optimal metastatic risk and worse regression (CI) or with optimal regression but worse metastatic risk (AI with CI). Baseline parameter values: *c*_1_ = *c*_2_ = *b*_2_ = *f*_2_ = 0.1; *λ* = 0.5; *k* = 0.4; *d*_1_ = 1.1; *d*_2_ = 0.95

As seen in Figure 3A, administering standard-of-care neoadjuvant AI treatment continuously for six months results in significant tumor regression (blue), which is improved by the addition of continuous administration of CI therapy (purple). However, the underlying phenotypic dynamics often represent very different outcomes even when tumor regression is similar. This is shown in Figure 3B, plotting the outcomes of phenotypic evolution on the CCR7–PD-L1 plane. The background is color-coded according to “metastatic risk” of each point in phenotype space (equation 5). The risk function is chosen such that *p*_1_ + *p*_2_ = 1 (here, *p*_1_ = *p*_2_ = 0.5), which has the convenient property that *m* ∈ [0,1] for every point in the simplex.

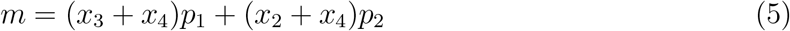

Importantly, this function increases with CCR7 expression and with PD-L1 expression. While the exact functional relationship between metastatic risk and expression of these markers is unknown, an increasing linear function is used as a first approximation, consistent with previous findings linking each expression axis to metastasis [37, 38, 39]. Therefore an ideal therapy would keep the CCR7 and PD-L1 expression low, residing in the top left corner for all of simulation time (*x*_1_ = 1) and giving a score of *m* = 0.

From these results it is apparent that there is a trade-off between tumor regression (Figure 3A) and metastatic risk (Figure 3B). Any therapy that results in both low CCR7 and PD-L1 expression will be associated with the lowest likelihood of developing metastatic disease. In Figure 3B, the CI treatment shows the best (lowest) metastatic potential score. This optimal solution comes with an important caveat: it is also associated with the worst tumor regression, second only to untreated growth (see Figure 3A). The highest tumor regression is associated with the continuous combination treatment (purple), which results in a relatively high metastatic risk in the top right corner of Figure 3B.

### 3.1. Phenotypic evolution under mono-and combination-therapy

In order to assess long-term outcomes, an “evolutionarily stable strategy” (ESS) can be directly calculated from the payoff table (Figure 2D), which indicates which phenotype will eventually dominant the tumor population. Each ESS shown shaded in Figure 4 holds for specific conditions (inequalities) [56]. Without treatment, the CCR7^−^ / PD-L1^+^ (Figure 4A, red) dominates if the cost of developing the PD-L1 strategy is less than immune cell kill rate in the absence of the immune escape mechanism (*c*_2_ < *k*_*T*_), otherwise the negative-negative phenotype wins. Under continuous AI treatment, the CCR7^+^ / PD-L1^+^ phenotype wins (Figure 4B, purple) if the cost of developing CCR7 expression is less than the kill rate due to AI drug therapy (*c*_1_ < *d*_1_) and the costs to CCR7/PD-L1 expression outweigh the cheating benefit of residing near a PD-L1^+^ cell in the presence of AI drug kill (−(*c*_1_ +*c*_2_) > *b*_2_ +*d*_2_*k*_*T*_ − *k*_*T*_) − *d*_1_). Under continuous CI treatment, the CCR7^−^ / PD-L1^−^ phenotype wins (Figure 4C, blue) if the costs to CCR7/PD-L1 expression are positive (*c*_1_ > 0; *c*_2_ > 0), otherwise neutral dynamics play out. Under continuous combination treatment, the CCR7^+^ / PD-L1^−^ phenotype wins (Figure 4D, yellow) if the cost to PD-L1 outweighs the sum of cost to AI resistance and the AI drug kill (*c*_2_ > *c*_1_ − 1). Interestingly, each of the four treatment scenarios results in a different dominant phenotype. This implies that the underlying phenotypic evolution may be controllable to any desired CCR7/PD-L1 expression by choosing which treatments to apply in sequence.

**Figure 4:**
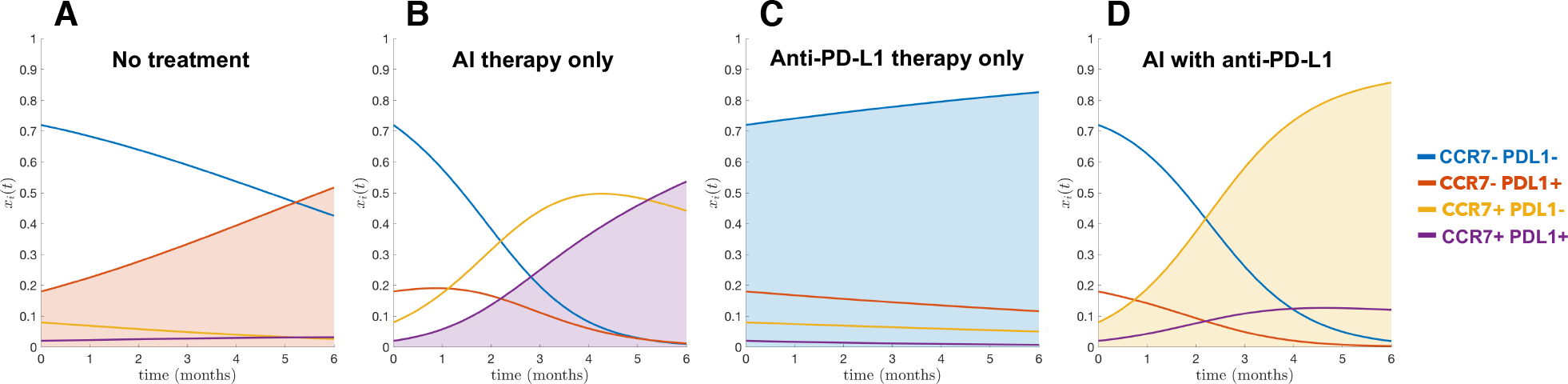
Evolutionary dynamics under continuous treatment. — model simulations are shown under no treatment (A), AI treatment (B), CI treatment (C), and both therapies (D). Relative tumor volume change is shown (black line, *υ*(*t*), top of each panel; equation 2) along with underlying phenotypic competitive dynamics (*x*_*i*_(*t*), bottom; equation 1). Interestingly, while each treatment shows varying tumor volume (with both AI treatments in B, D showing tumor decrease), the emerging dominant phenotype for each treatment is different. (*c*_1_ = *c*_2_ = *b*_2_ = *f*_2_ = 0.1; *λ* = 0.5; *k* = 0.4; *d*_1_ = 1.1; *d*_2_ = 0.95)

### 3.2. Transient dynamics: a virtual PEPI score

In order to fully explore the trade-off between tumor regression and metastatic risk, it is useful to visualize these dueling axes of the trade-off between metastatic risk and tumor regression, as shown in Figure 5. All therapies begin with an identical initial condition (black dot) and track tumor volume (vertical axis) and time-weighted average metastatic risk (horizontal axis) for 6 months of neoadjuvant therapy. Here, the optimal therapy would direct the curve towards the origin.

The model’s true utility comes when describing the transient dynamics of the underlying PEPI indicators. In Figure 5A, the trajectory of monotherapies and combination therapies is shown for no treatment (black) and 5 different treatment regimens. Intermittent CI schedules were not analyzed due to anti-PD-L1 having a long effective half-life in patients. The optimal metastatic risk score (horizontal axis) is associated with CI monotherapy (red dot) but this is the second worst tumor regression (vertical axis). The current standard of care (AI; blue dot) decreases tumor volume compared to no treatment (black dot) but is the worst outcome for metastatic risk. Continuous AI and CI therapy results in a better overall outcome than the standard of care by lowering *both* tumor regression and metastatic risk.

Delayed treatment strategies are tested in Figure 5B. The legend shows the delay time (months) of one drug with respect to continuous (six month) administration of the other: the zero entry indicates continuous treatment of both. The model predicts that the global optimum tumor regression is achieved through continuous CI therapy coupled with one month delayed AI. This results in a temporary increase in tumor volume for the first month. However, fast regression for the final five months of neoadjuvant therapy leads to the smallest tumor burden at time of surgery. These results are relatively robust to changes in parameters, provided the inequalities described in section 3.1 remain true. These inequalities determine the long-term dominant phenotypes, and variations in parameters show that continuous CI and 1 month delayed AI remains optimal in most circumstances (see Supplemental Table 1, Figure A1), except for the cases of large costs or low initial expression of CCR7 or PD-L1.

**Figure 5:**
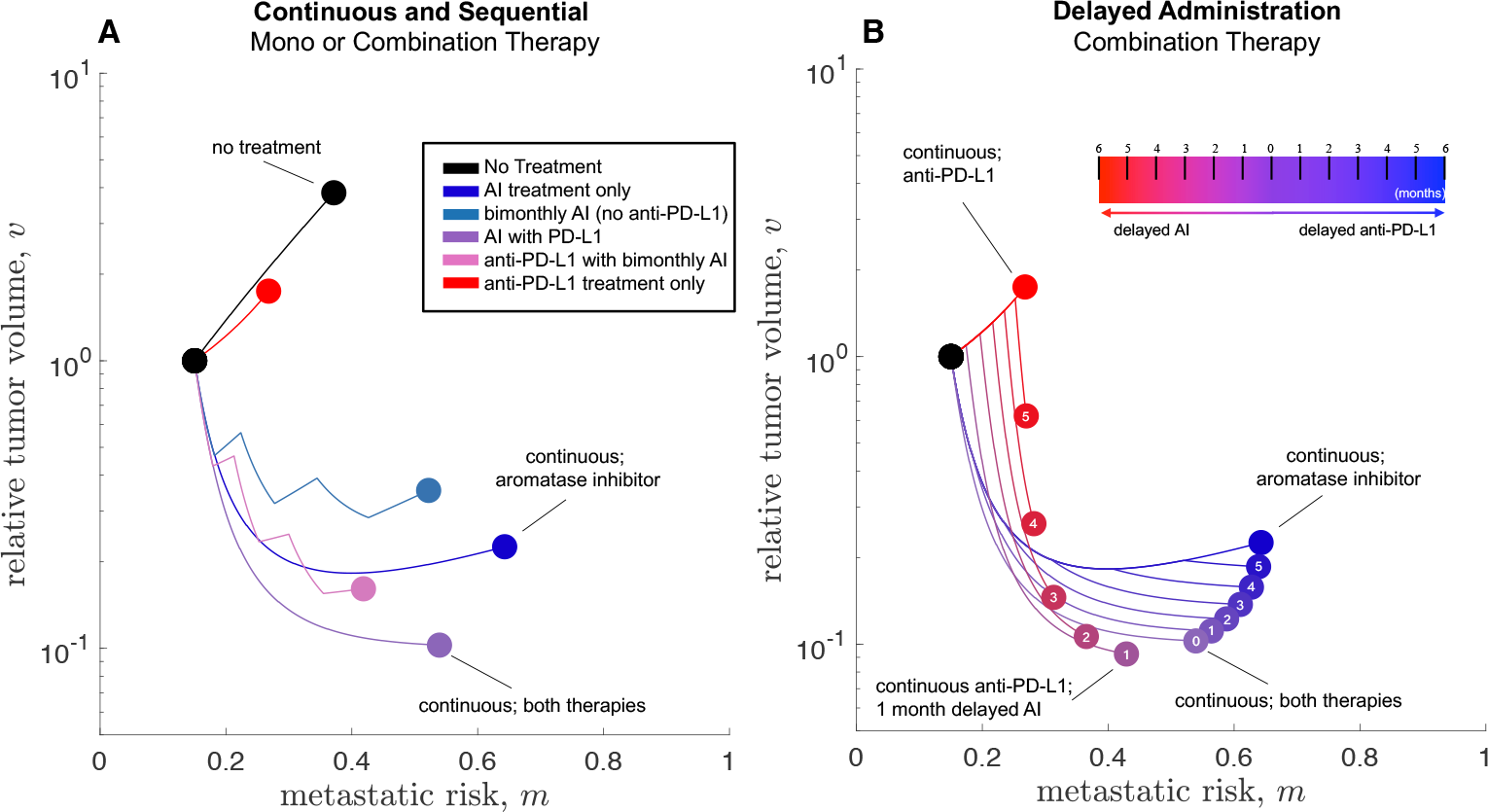
Mono-and combination therapies. — The trajectory through metastatic risk space (A), change in tumor burden (B) and total score (C) is shown for 8 therapeutic strategies (see legend). Strategies are loosely grouped into: 1) continuous AI, 2) continuous CI, and 3) alternating/synchronized sequences. Parameter values: *c*_1_ = *c*_2_ = *b*_2_ = *f*_2_ = 0.1; *λ* = 0.5; *k* = 0.4; *d*_1_ = 1.1; *d*_2_ = 0.95

## 4. Discussion

The PEPI score is used to predict prognosis after neoadjuvant therapy, and combines tumor size, Ki-67 proliferation index, hormone status, and lymph-node involvement. The PEPI score aims to optimize all four, yet gives no information about transient scores, or which of the four factors to prioritize. The purpose of this study is to highlight the inherent trade-off between two of these indicators: metastatic risk and tumor regression. Our results suggest that one month of delayed aromatase inhibitors combined with continuous checkpoint inhibitors leads to optimal tumor regression. However we can imagine scenarios where other factors might be prioritized. A patient with an large, aggressive and invasive tumor may opt to receive a stronger dose of combination therapy, lowering the tumor volume (vertical axis, Figure 3C) at the expense of increased metastatic risk (lateral axis, Figure 3C). On the other hand, since neoadjuvant therapy is followed by surgery at the end of the 6 month therapy. This may indicate that in some instances it is wise to optimize lowering a patient’s metastatic risk rather than decreasing tumor burden. In other scenarios, surgery may not be plausible. The two axes need to be considered concurrently as they are coupled; a higher metastatic score with a small tumor is likely less dangerous than the same with a large tumor at the end of neoadjuvant therapy. Interestingly, using a simple minimization of distance to the origin (Figure 5) resulted in the same optimal treatment of 1 month delayed AI under a variety of parameters tested (see Table 1).

We are often inclined to define the success of a therapy as the eradication of the tumor. Neoadjuvant therapies are followed by surgical removal of the primary tumor, altering the desired treatment outcome. Clinicians are tasked with creating or maintaining an operable tumor volume with the primary goal of delaying or even preventing post-surgical recurrence. This setting allows for more nuanced approaches. We no longer have to maximize tumor cell kill at the risk of driving drug resistance or increasing metastatic potential. We can begin to optimize over metrics such as long-lasting systemic immune memory and selection for less aggressive phenotypes if recurrence does arise. As treatment options become more diverse and more sophisticated so too will our applications of such treatments, and therefore the need for applying more integrated approaches like the one we develop here.

Optimizing multiple output variables (regression and metastasis) is not a straightforward task, but mathematical models are a powerful abstraction of clinical intuition, enabling hypothesis generation and testing of clinical assumptions. Especially when developed in collaboration with clinicians, such models provide clarity and power despite their simplicity and built-in assumptions, emphasizing their ability to define novel therapeutic regimens. Here we have found that combination therapy is likely to be better than either monotherapy for reducing both tumor burden and metastatic risk. Furthermore introducing a one month delay for AI can strengthen the outcome.

## Supplemental Material

**Table 1:** Optimal Tumor Regression. Each ordered pair represents the optimal strategy (Months delayed AI, Months delayed anti-PD-L1), such that (0,0) represents no delay, continuous treatment of both. Here optimum is measured minimum tumor burden: min(*υ*) at the end of neoadjuvant therapy.This table represents outcomes for Figure A1: baseline parameters are *b*_2_ = 0.1; *c*_1_ = 0.1; *c*_2_ = 0.1; *f*_1_ = 0.1; *f*_1_ = 0.1; *k*_*T*_ = 0.4; *λ* = 0.5; *d*_1_ = 1.1; *d*_2_ = 0.95, unless otherwise noted. Most common optimal strategy (1,0) shown in red.

| Parameter Value | 0.05 | 0.15 | 0.25 | 0.35 |
| --- | --- | --- | --- | --- |
| Varied $b_2$ | (1,0) | (1,0) | (1,0) | (1,0) |
| Varied $c_1$ | (1,0) | (1,0) | (0,0) | (0,0) |
| Varied $c_2$ | (1,0) | (1,0) | (0,0) | (0,0) |
| Varied $f_1$ | (0,0) | (1,0) | (1,0) | (1,0) |
| Varied $f_2$ | (0,0) | (1,0) | (1,0) | (1,0) |

**Table 2.** Optimal Tumor Regression and Metastasis. Each ordered pair represents the optimal strategy (Months delayed AI, Months delayed anti-PD-L1), such that (0,0)represents no delay, continuous treatment of both. Here optimum is measured as distance from the origin: 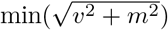 at the end of neoadjuvant therapy. This table represents outcomes for Figure A1: baseline parameters are *b*_2_ = 0.1; *c*_1_ = 0.1; *c*_2_ = 0.1; *f*_1_ = 0.1; *f*_1_ = 0.1; *k*_*T*_ = 0.4; *λ* = 0.5; *d*_1_ = 1.1; *d*_2_ = 0.95, unless otherwise noted. Most common optimal strategy (1,0) shown in red.

| Parameter Value | 0.05 | 0.15 | 0.25 | 0.35 |
| --- | --- | --- | --- | --- |
| Varied $b_2$ | (1,0) | (1,0) | (1,0) | (1,0) |
| Varied $c_1$ | (1,0) | (1,0) | (1,0) | (0,0) |
| Varied $c_2$ | (1,0) | (1,0) | (1,0) | (1,0) |
| Varied $f_1$ | (1,0) | (1,0) | (2,0) | (2,0) |
| Varied $f_2$ | (1,0) | (1,0) | (2,0) | (2,0) |

**Figure 1:**
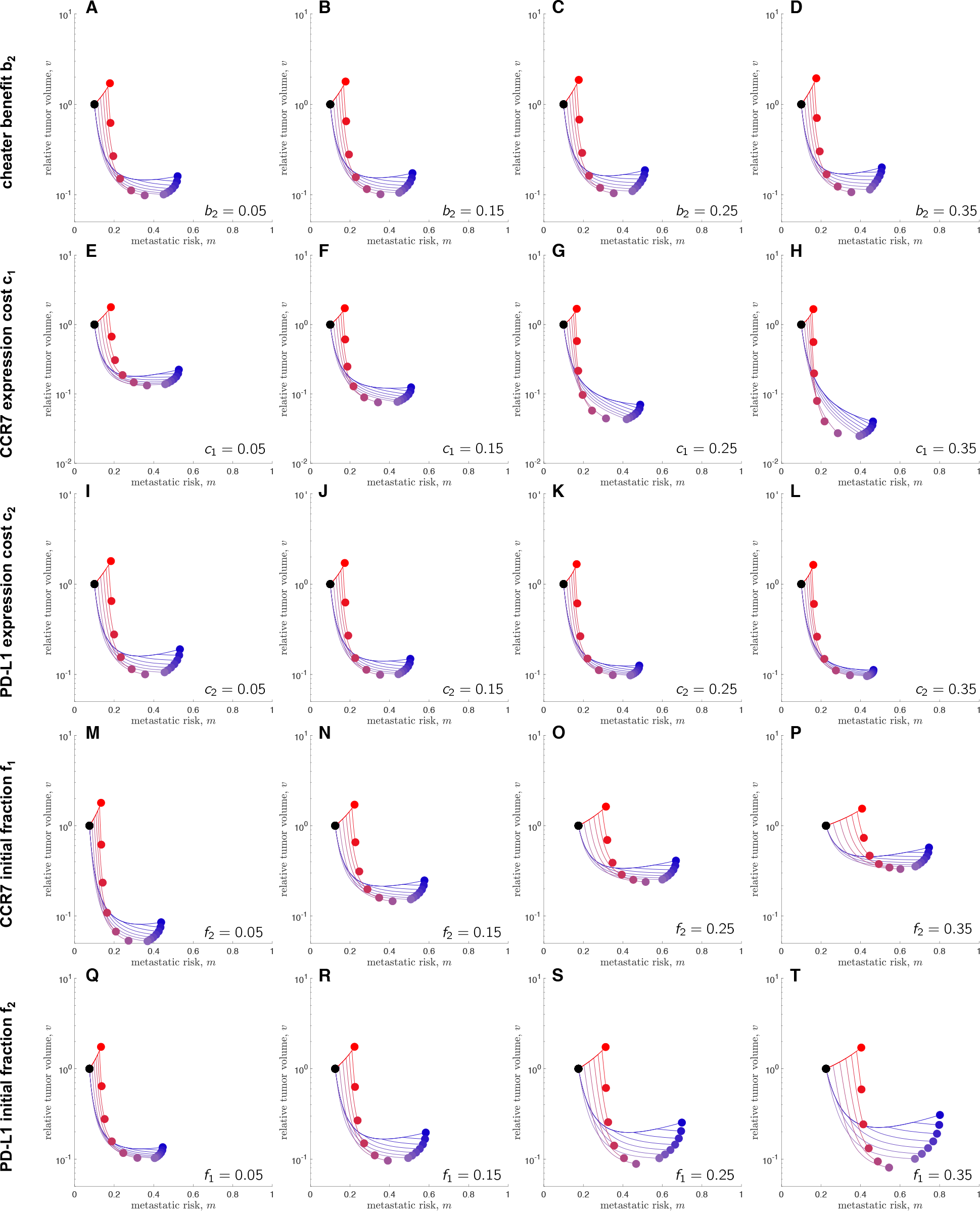
Parameter sensitivity analysis. — Baseline parameters are *b*_2_ = 0.1;*c*_1_ = 0.1;*c*_2_ = 0.1;*f*_1_ = 0.1;*f*_1_ = 0.1;*k*_*T*_ = 0.4;*λ* = 0.5;*d*_1_ = 1.1;*d*_2_ = 0.95, unless otherwise noted.

